# Hippocampal subfields support human novelty detection via distinct signals

**DOI:** 10.64898/2026.09.08.748144

**Authors:** Jörn A. Quent, Kaixiang Zhuang, Xinyu Liang, Debin Zeng, Yun Wang, Jianfeng Feng, Jordan DeKraker, Boris C. Bernhardt, Deniz Vatansever

## Abstract

Detecting novelty is fundamental to adaptive memory, yet the neural comparisons that allow new events to be distinguished from prior experience are not fully understood. Hippocampal novelty signals are often framed as associative comparator responses that register violations of learned relationships. However, recognition memory models also predict a broader form of comparison: a global, item-based mismatch between an incoming stimulus and aggregate memory traces. Whether such global matching is implemented within human hippocampal circuitry, and whether it contributes to memory decisions, remains unknown. Using ultra-high-field 7T fMRI during dense sampling of a continuous object recognition task (76,800 trials across five sessions), we estimated both univariate novelty responses and multivariate global mismatch signals across hippocampal subfields. CA1 carried the novelty response that most reliably predicted accurate novelty detection, whereas the subiculum expressed a global mismatch signal that predicted detection independently of CA1. This subicular signal also shaped subsequent recognition behaviour, in which low global mismatch was associated with false alarms to novel objects that propagated into later hits. Finally, subicular global mismatch scaled with the experienced similarity of past events, consistent with an item-based comparison against accumulated memory. These findings identify distinct hippocampal mismatch signals for novelty detection and extend the human hippocampal comparator role beyond local associative mismatch to include global, item-based matching.

## Introduction

The ability to detect novel information is a foundational operation of our adaptive memory system. Constrained by finite attentional and encoding resources, the human brain preferentially commits unexpected events into memory, thereby improving the efficiency in guiding subsequent goal-directed behaviour (*1, 2*). Importantly, this prioritisation is more than an epiphenomenon of everyday learning. The same novelty signals can be harnessed as interventional tools to strengthen memory through novelty-induced behavioural tagging or consolidation (*3*–*6*), and their degradation can mark the early stages of neurodegenerative disease (*7*). Therefore, deciphering the computational and neural mechanisms by which the human brain computes novelty remains central both to theories of memory and to their translational potential.

Convergent clinical and neuroimaging evidence places the hippocampus at the centre of novelty detection. Alongside its canonical role in declarative memory (*8, 9*), it is reliably engaged by novel stimuli across species and recording modalities, including rodent electrophysiology (*10, 11*), and human single-unit (*12*), intracranial (*13*) and fMRI recordings (*14–16*). However, novelty sensitivity is not uniform across the hippocampal circuitry. For example, hippocampal subfields perform specialised yet complementary computations (*17–19*), with the dentate gyrus (DG) and Cornu Ammonis 3 (CA3) dynamics biased towards pattern separation, while the CA3-CA1 dynamics support pattern completion (*20, 21*).

Within this canonical circuit, CA1 is anatomically well positioned to act as a comparator for novelty detection (*22*). CA1 receives direct sensory projection from the entorhinal cortex, and indirect, mnemonically transformed input via the DG and CA3, enabling a comparison between current input and stored predictions (*11, 23*). Predictive-coding accounts recast this computation as a prediction error, arising when higher levels of a memory hierarchy fail to anticipate incoming information (*24–27*). Consistent with this view, CA1 responses scale parametrically with the degree of mismatch between expectation and observation. For instance, re-arranging learned temporal sequences of objects (*28*) or altering the layout of learned virtual environments (*23*) elicits graded CA1 activity that tracks the strength of the violated prediction. A crucial caveat, however, is that in nearly all of this prior work, mismatch has been operationalised as relational violations of learned item-to-item or item-to-context associations (*23, 28–30*), with recent evidence indicating that the hippocampal comparator is engaged specifically by mismatch with episodic memories (*31*) and that particular subfields are contribute when these associations are established and well-learned (*22*). Thus, beyond relational comparison, what remains unclear is whether the human hippocampus also contributes to a more fundamental, non-associative form of mismatch.

Classical models of recognition memory formalise the old/new decision as a process of global matching, in which a probe is compared in parallel against all stored memory traces, generating a continuous signal that scales with the probe’s aggregate similarity to accrued experience (*32–34*). Low values index a global mismatch, providing a continuous, non-associative novelty signal that is distinct from the relational mismatch, and is computed over context-specific violations of learned associations. Although global matching has largely been attributed to peripheral regions in the medial temporal lobe (*35, 36*), the hippocampus nonetheless also responds robustly to the novelty of individual items in standard recognition paradigms in which no learned association is violated (*15, 20, 37*). Recent computational work further shows that representational similarity between stimuli systematically modulates novelty signals and novelty-driven behaviour (*38*), underscoring that this item-to-memory similarity is intrinsic to the computations of novelty.

Collectively, these observations motivate two frameworks with opposing, yet testable predictions. If the hippocampal comparator is intrinsically associative, then item-based global matching should only be supported by regions outside the hippocampus and thus should not involve any subfield (*34, 39*). Instead, if the hippocampus implements a general, similarity-based comparison, then at least one subfield should carry a graded global mismatch signal even when no relational structure is violated. Since a univariate novelty response and a multivariate matching computation need not be co-localised, the two may dissociate across subfields, such that the region generating the largest novelty response is not necessarily the one that computes global mismatch. For instance, as the principal output stage of the hippocampus, the subiculum is an a priori candidate for this global comparator role (*40*). The subiculum pools convergent input from many CA1 neurons (*41*) and encodes predictive representations over long time scales (*42*), providing a suitable opportunity to compare an item against aggregate experience instead of any specific association. At a broader level, adjudicating between these alternatives determines whether the hippocampal comparator is solely relational or additionally computes the item-to-memory match that recognition theory attributes to familiarity.

In addition to the established univariate novelty response, here we tested whether, and in which hippocampal subfield the human brain computes a global, item-based mismatch signal. We made three predictions: We reasoned that (i) if the hippocampal comparator is not confined to a relational structure, at least one subfield should express a multivariate global mismatch signal that predicts novelty detection over and above the univariate novelty response; (ii) if this signal implements global matching, it should also support mnemonic decisions, such that early false alarms propagate into later hits; and (iii) if it reflects a prediction error whose magnitude is shaped by experience, it should scale with an item’s aggregate similarity to a participant’s past, and this influence should decay with time. We further examined between-participant variability in how this signal influenced novelty decisions, assessing whether response bias and memory sensitivity modulated its impact. In order to test these predictions, we acquired a large-scale dataset in which twenty participants judged whether a naturalistic object image drawn from the THINGS database (*43*) was old or new, across 76,800 trials (3,840 per participant) distributed over five separate days. This dense-sampling design ensured a continuous supply of genuinely novel items while allowing similarity and lag to vary naturally.

Combining ultra-high-field 7T fMRI, automated hippocampal subfield segmentation (*44, 45*) and Bayesian hierarchical modelling, we found that CA1’s univariate novelty response most reliably predicted accurate detection, although the response magnitude itself was greatest in CA3. By contrast, the subiculum showed no robust univariate novelty response but expressed a multivariate global mismatch signal that predicted novelty detection independently of CA1. This global, item-based mismatch signal propagated false alarms into subsequent hits and scaled with experienced semantic and visual similarity in a time-dependent manner. Contrary to prevailing accounts, these results identify a non-associative, item-based comparison as a hippocampus-based computation and motivate extending the comparator framework to include global mismatch, with hippocampal subfields computing prediction errors at different granularities, from the local, relational mismatch attributed to CA1 to global mismatch in the subiculum. More broadly, these findings may provide a principled bridge between hippocampal circuit mechanisms significantly furthering our understanding of hippocampal function.

## Results

### Individual items are reliably recognised during continuous object recognition

Across five fMRI sessions, participants (n = 20, 14 females, mean age = 24.56 years) completed 60 runs of a continuous object recognition task in which naturalistic object images were presented every 4 s, interleaved with blank events (64 image trials per run; Fig. 1A). Each image was shown three times across the experiment, and on each trial, participants judged whether it was new (never previously seen) or old (seen in the current or a previous session; Fig. 1B). Because images recurred, the proportion of new items steadily decreased while the number of old items in memory increased, creating a regime in which any putative global-matching signal would accumulate against an expanding memory set and enabling calculation of both univariate novelty and multivariate mismatch signals (Fig. 1C).

**Fig. 1.**
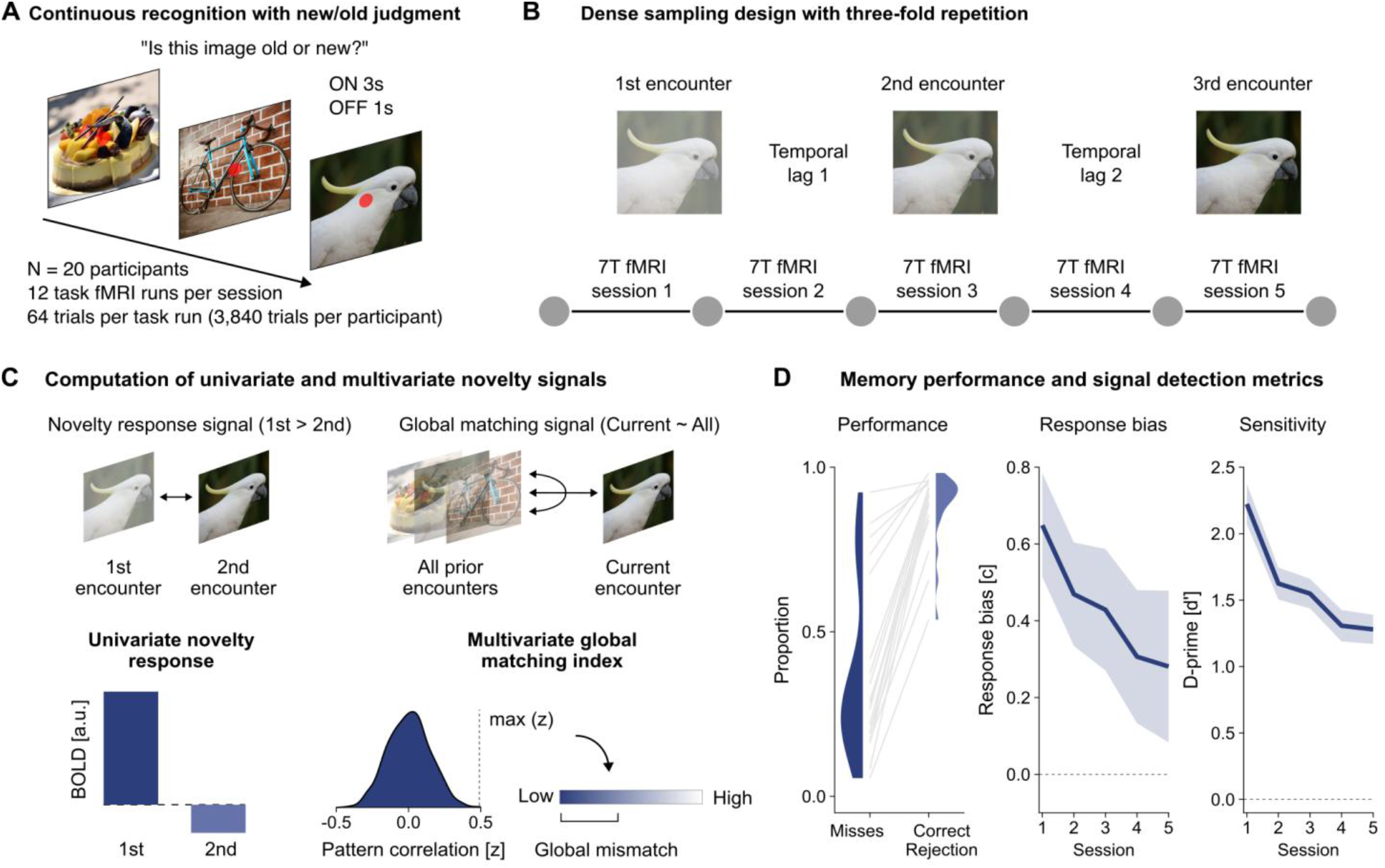
Continuous recognition task and behavioural performance. **(A)** Continuous recognition with old/new judgements. Naturalistic object images from the THINGS database were presented every 4 s, interleaved with blank events, in 12 runs with 64 image trials per sess ion. On each trial, participants made a speeded old/new judgement, indicating whether the image was new (never previously seen) or old (seen during the current or an earlier session). **(B)** Dense sampling design with three-fold repetition. The full experiment spanned across five separate scanning sessions on five separate days, over the course of which each image was presented three times in total. The temporal lag between encounters of the same image varied between seconds and days. **(C)** Computation of univariate and multivariate novelty signals. We computed and contrasted two novelty signals, which support novelty detection. First, we calculated a univariate novelty response by subtracting the estimated average single trial beta for the 2nd encounter from the 1st encounter. Second, we derived a multivariate global mismatch signal by calculating the Fisher-z transformed Pearson pattern correlation between the current stimulus and all stimuli previously encountered and defining our index as the maximum of that distribution. To avoid contamination with temporal autocorrelation, same-run trials were excluded from this analysis. **(D)** Memory performance and signal detection metrics. Left: Misses and correct-rejection rates per participant; correct-rejection rates exceeded miss rates for every participant (grey lines). Their distributions are summarised as half-violine plots. Middle: Session-level response bias (c) showing no net group-level change. Right: Session-wise sensitivity (d′), which declined over time but remained well above chance in the final session. Lines with ribbon illustrate group-level mean ± standard error of the mean (SEM).

We first confirmed that the employed rapid event-related procedure sustained participant engagement in this dense-sampling design, containing a large number of trials, runs and sessions. Providing no indication of fatigue or waning motivation, response rates remained consistently high and did not decline over the length of the experiment (session-level rates 82–100%, β = 0.061, 95% CI [−0.46, 0.58]; Fig. S1A), and reaction times were likewise stable (β = 0.013, 95% CI [−0.13, 0.15]), averaging 1.1 s across sessions (SD = 0.19; Fig. S1B). Despite the large and growing memory load, participants discriminated old from new images with high efficiency, with correct-rejection rates exceeding miss rates for every participant, tested via a Bayesian *t*-test, d = 2.3, BF_10_ > 1000; session-wise d = 1.9– 2.4 (Fig. 1D). As expected, sensitivity (d′) declined over time (β = −0.94, 95% CI [−1.3, −0.54]; Fig. 1D), yet remained well above chance even in the final session (BF_10_ > 1000, mean = 1.3, SD = 0.49). Response bias (c) showed no net group change (β = −0.34, 95% CI [−0.75, 0.08]; Fig. 2B). While a few participants gradually became more conservative, most became slightly more liberal, possibly tracking the rising proportion of old images (range −0.8 to 0.18). Together, these behavioural results provide a reliable, high-quality dataset to interrogate novelty signals in the human hippocampus.

**Fig. 2.**
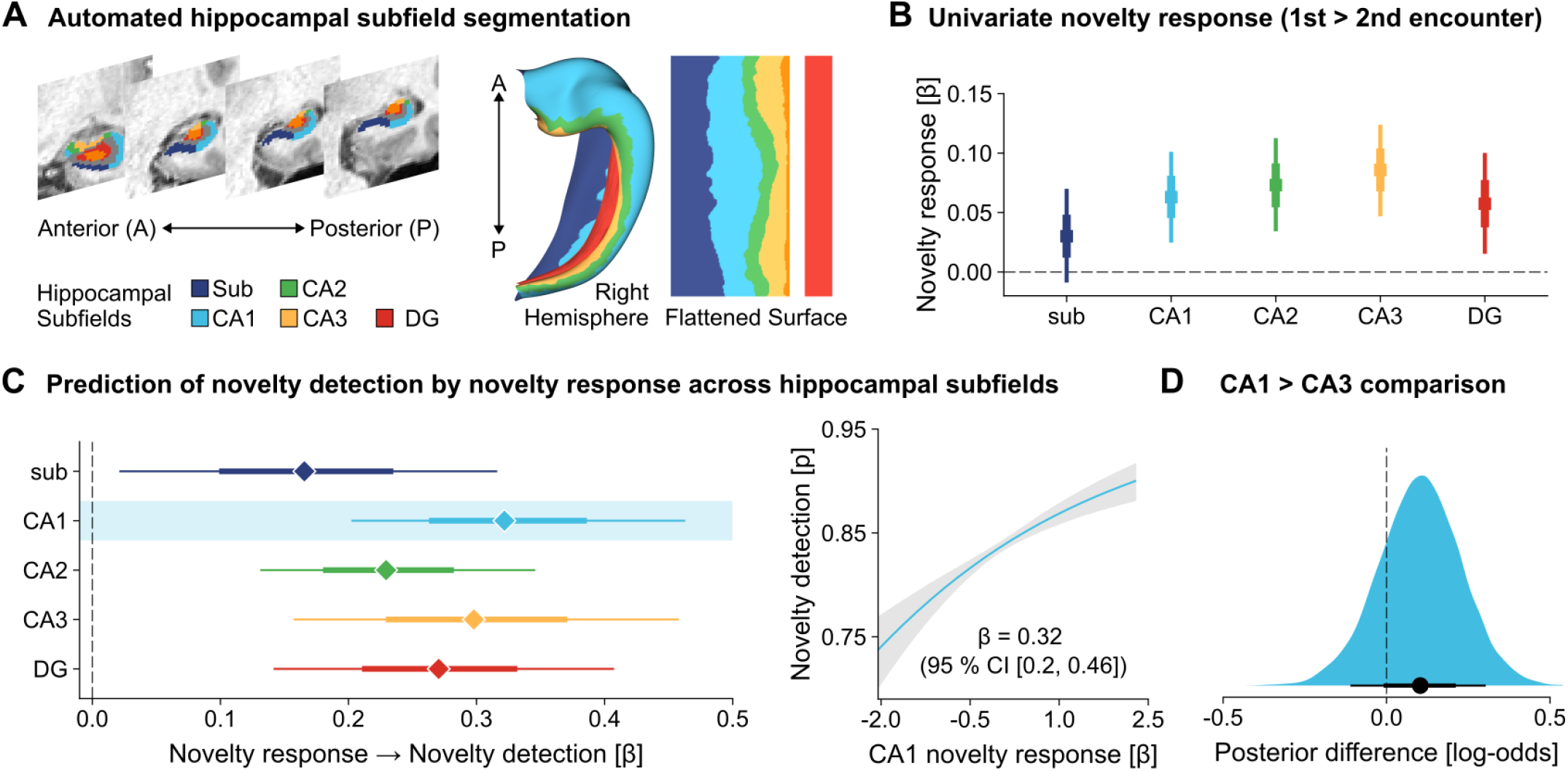
The univariate novelty response and its prediction of detection across hippocampal subfields. **(A)** Anatomical schematic of the automatically segmented hippocampal subfields. Four representative (coronal) slices were selected to illustrate segmentation success of a single participant (right hemisphere). Subfields are additionally illustrated on the canonical HippoMaps (45) surface. **(B)** Univariate novelty response. Item-level univariate novelty response (1st minus 2nd encounter) for each subfield (subiculum, CA1, CA2, CA3 and DG), estimated with Bayesian hierarchical models. Robust responses were present in all subfields except the subiculum, peaking in CA3. Posterior distributions are summarised via their median (centre square) and the 66% and 95% CI. **(C)** Prediction of novelty detection by novelty response across hippocampal subfields. The novelty response in CA1 most strongly predicted novelty detection. Posterior distributions are summarised via their median (centre diamond) and the 66% and 95% CI. To illustrate the relationship between CA1’s novelty response and novelty detection, we show marginal the prediction averaged across participants and co-variates (shaded band = 95 % CI). **(D)** Direct posterior contrast β (CA1) − β (CA3) testing the CA1-versus-CA3 dissociation in the univariate novelty response. The posterior difference distribution is summarised via the median (centre dot) and the 63% and 90% CI.

### CA1’s univariate novelty response most reliably predicts accurate detection

Following prior work, our initial objective was to characterise the univariate hippocampal novelty response, defined at the item-level as the difference between an image’s first and second encounter (1st > 2nd) and averaged within each subfield (Fig. 2A). The third encounter was deliberately excluded to avoid contamination by reactivation-related processes. All analyses used Bayesian hierarchical models with full participant-level random effects.

The results revealed robust univariate novelty responses, widespread across the hippocampal subfields (Fig. 2B). The response peaked in CA3 (β = 0.09, 95 % CI [0.05, 0.12]), followed by CA2 (β = 0.07, 95 % CI [0.03, 0.11]), CA1 (β = 0.06, 95 % CI [0.02, 0.10]) and the DG (β = 0.06, 95 % CI [0.02, 0.10]), while only the subiculum lacked a robust univariate response (β = 0.03, 95 % CI [−0.01, 0.07]). Since univariate novelty responses can reflect extraneous processes other than novelty detection (e.g. attention, encoding, and arousal), we next tested whether response magnitude predicted trial-wise detection success using logistic models.

Responses in every subfield predicted detection success, but the association was strongest in CA1 (β = 0.32, 95 % CI [0.20, 0.46]; Fig. 2C), closely followed by CA3 (β = 0.30, 95 % CI [0.16, 0.46]), the DG (β = 0.27, 95 % CI [0.14, 0.41]), CA2 (β = 0.23, 95 % CI [0.13, 0.35]) and the subiculum (β = 0.17, 95 % CI [0.021, 0.32]). When entered into a joint model and directly contrasted via Bayesian hypothesis testing, CA1’s association with novelty detection remained robust, while CA3’s did not, with moderate evidence for a difference between the subfields (0.1, 90% CI [−0.11, 0.30], BF_10_ = 3.87; one-sided; Fig. 2D). Thus, although CA3 exhibited the largest novelty response, CA1 most reliably tracked successful novelty detection, consistent with its canonical role as the circuit’s comparator readout.

### A subicular global mismatch signal predicts novelty detection independently of CA1

Having established that CA1’s univariate novelty response most reliably predicted detection, we next tested our first prediction, that hippocampal subfields additionally implement the parallel item-to-memory comparison posited by global-matching models. For each trial, we computed pairwise correlations between the current multivariate activity pattern and the patterns evoked on all previous trials in a given subfield (excluding same-run pairs), and defined a global matching index as the maximum of these correlations (Fig. 1C). Throughout, negative coefficients indicate that greater mismatch was associated with higher outcome values, whereas positive coefficients indicate the inverse. Since Pearson correlation is scale-free, this index is insensitive to uniform activation differences and is therefore dissociable from the univariate novelty response.

We identified a strong global mismatch signal exclusively in the subiculum that predicted accurate novelty detection across participants (β = −0.26, 95% CI [−0.41, −0.11]; Fig. 3A & 3B). In contrast to the univariate analyses, no other subfield showed a robust effect, including CA1 (CA3: β = −0.12, 95% CI [−0.26, 0.02]; DG: β = −0.08, 95% CI [−0.21, 0.04]; CA1: β = −0.07, 95% CI [−0.21, 0.07]; CA2: β = −0.04, 95% CI [−0.20, 0.15]). The subicular signal also tracked decision speed, a widely used proxy for decision confidence (*46*). Shorter (log) reaction times predicted accurate detection (β = −0.55, 95% CI [−0.96, −0.16]; Fig. S2), and in parallel a lower subicular matching index (greater mismatch) predicted faster responses (β = 0.037, 95 % CI [0.022, 0.054]; Fig. 3B), providing converging evidence that a global-matching computation in the subiculum supports robust novelty detection.

**Fig. 3.**
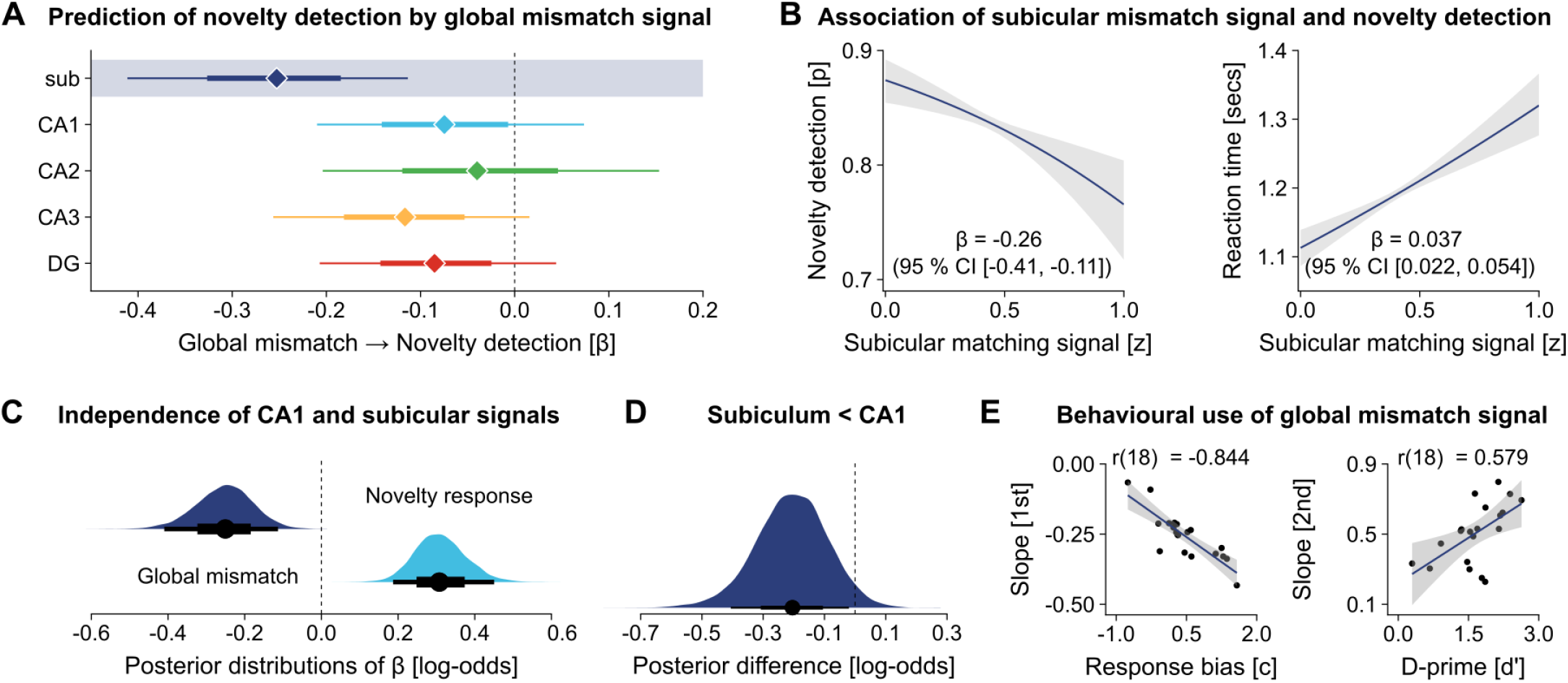
A global mismatch signal in the subiculum predicts novelty detection independently of CA1. **(A)** Prediction of accurate novelty detection from the global matching index in each subfield. Only the subiculum displayed a robust (negative) effect. Posterior distributions are summarised via their median (centre diamond) and the 66% and 95% CI. **(B)** Association of subicular mismatch signal and novelty detection. The subicular mismatch signal predicted novelty detection success as well as more confident (i.e. faster) responses. Marginal predictions are visualised averaged across participants and co-variates (shaded band = 95 % CI). **(C)** Joint model containing both CA1’s univariate novelty response and the subicular global mismatch signal. Each retained its full predictive strength, indicating two separable contributions to novelty detection. Posterior distributions are summarised via their median (centre dot) and the 66% and 95% CI. **(D)** Direct posterior contrast β (subiculum) − β (CA1) testing the multivariate global-mismatch dissociation. The posterior difference distribution is summarised via the median (centre dot) and the 63% and 90% CI. **(E)** Behavioural use of global mismatch signal. Left: Slopes based on novelty detection prediction plotted against response bias (c) showing more conservative participants were more strongly guided by the global mismatch signal. Right: Slopes based on retrieval success prediction plotted against sensitivity (d′): better-performing participants showed tighter coupling between the matching signal and retrieval success on the second presentation. Points denote participants; shaded bands = 95% CI.

Three control analyses further validate the subicular mismatch signal’s contribution to novelty detection. First, because the matching index is defined to be independent of overall activation, we entered CA1’s univariate novelty response and the subicular mismatch signal into a single joint model, in which both retained their full predictive strength (CA1 novelty response β = 0.31, 95% CI [0.19, 0.45]; subicular mismatch β = −0.25, 95% CI [−0.41, −0.11]; Fig. 3C). This pattern suggests two separable contributions to novelty detection: (1) a local, univariate comparator centred on CA1 and (2) a global, multivariate computation in the subiculum. Second, we tested if global matching in the subiculum was more strongly associated with novelty detection compared to global matching in CA1 in another joint model using Bayesian hypothesis testing. We found strong evidence that that the association is stronger in the subiculum (−0.21, 90% CI [−0.41, −0.02], BF_10_ = 28.76; one-sided; Fig. 3D). Third, although image memorability, nameability, and recognisability each predicted behaviour, none predicted the subicular index (Fig. S3), arguing against these stimulus-driven factors as potential confounds.

We next tested our second prediction seeking a behavioural signature that would distinguish global matching process from a simple correlation. If the subiculum implements the comparator that underlies recognition decisions, then a high global match, which causes a novel image to be falsely endorsed as old, should propagate early false alarms into subsequent hits when that image genuinely recurs. Consistent with this prediction, the first-encounter matching signal positively predicted subsequent hits for both the second (β = 0.51, 95% CI [0.36, 0.66]; Fig. S4) and the third encounter (β = 0.44, 95% CI [0.26, 0.62]; Fig. S4), which represents a feature of a global-matching comparator that a reverse-causality account cannot accommodate.

In a final exploratory analysis, we additionally tested whether the identified global mismatch signal’s behavioural impact varied across individuals by extracting participant-specific slopes from the hierarchical models and correlated them with signal-detection measures using Bayesian Pearson correlations. Participants whose novelty judgements were more strongly influenced by subicular global mismatch tended to adopt a more conservative response criterion (c), r(18) = −0.844, BF_10_ > 1000. (Fig. 3E). In addition, higher-performing participants (higher d′) showed tighter coupling between the matching signal and retrieval success on the second encounter, r(18) = 0.579, BF_10_ = 7.78.

Together, these results suggest that subicular global mismatch supports novelty detection beyond univariate and multivariate contributions of CA1. Furthermore, global mismatch is a consistent feature of novelty detection, which predicts response confidence and in line with theoretical predictions also consistently relates to subsequent memory. Finally, its behavioural expression varies with both decision criterion and memory quality.

### The subicular global mismatch signal scales with the similarity of prior experience

In our third and final prediction, we reasoned that if the subicular signal computes a global mismatch whose magnitude was shaped by experience, then it should scale with the extent to which each probe resembles a participant’s prior experience and should impair novelty detection accordingly. We therefore quantified image-to-image similarity using the 66 embedding dimensions derived from large-scale human similarity judgements (*47*), applying the same maximum-similarity logic used for the neural index. We further classified each dimension as semantic or visual (*48*) (Table S1; Fig. 4A) to estimate and contrast their contributions.

**Fig. 4.**
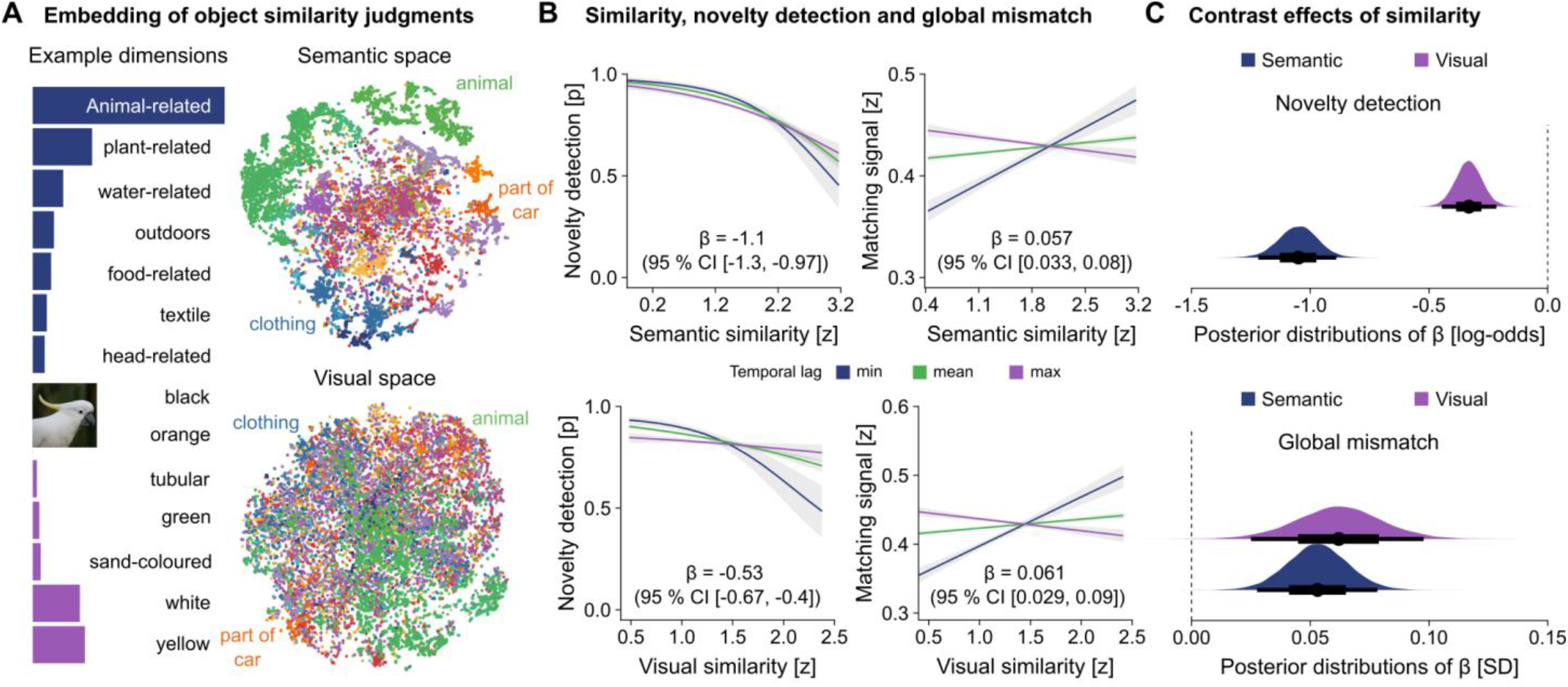
Subicular global mismatch signal scales with semantic and visual similarity to past experience. **(A)** Classification of the 66 human-similarity embedding dimensions into semantic and visual components. We show how an example image (cockatoo_04s.jpg) varies along these dimensions, which are illustrated by seven representative dimensions each. Due to copyright reasons, the displayed image has been replaced with a similar CC0 image from THINGSplus for illustrative purposes. Embeddings based on these dimensions are visualised as 2D t-SNE maps. Each dot represents an experimental stimulus coloured by its broadest category (e.g. animal). **(B)** Marginal effects of semantic and visual similarity on novelty detection (left) and on the neural global matching index (right): higher similarity impaired detection and reduced global mismatch, with both effects attenuating at longer lags since the most similar prior image. The interaction between similarity and temporal lag is visualised by separate slopes for the minimum, mean and maximum temporal lag. Marginal predictions are visualised averaged across participants and co-variates (shaded band = 95 % CI). **(C)** Contrast effects of similarity. Joint models entering semantic and visual similarity together; each retains independent predictive strength for both behaviour (top) and the neural signal (bottom). Posterior distributions are summarised via their median (centre dot) and the 66% and 95% CI.

As expected, high semantic (β = −1.1, 95% CI [−1.3, −0.97]) and visual similarity (β = −0.53, 95% CI [−0.66, −0.40]) each reduced the probability of correct novelty detection (Fig. 4B), with a larger impairment for semantic than visual similarity. In both cases, this interference was modulated by the (log) lag since the most similar prior image (semantic: β = 0.32, 95% CI [0.082, 0.58]; visual: β = 0.45, 95% CI [0.16, 0.76]), indicating that recently encountered similar images strongly impaired novelty detection. This interference decayed over time, more steeply for visual than for semantic similarity, consistent with more durable semantic predictions.

The neural signal mirrored this behavioural pattern. Greater similarity reduced global mismatch for both semantic (β = 0.057, 95% CI [0.033, 0.08]) and visual dimensions (β = 0.06, 95% CI [0.028, 0.089]; Fig. 4B), and these effects weakened at longer lags (semantic: β = −0.14, 95% CI [−0.19, −0.094]; visual: β = −0.18, 95% CI [−0.22, −0.13]). Equivalent results held when similarity was instead defined from a pre-trained CLIP model (Fig. S5). Modelling semantic and visual similarity jointly preserved their robust prediction of both behaviour (semantic β = −1.1, 95% CI [−1.2, −0.89]; visual β = −0.33, 95% CI [−0.44, −0.22]) and the neural signal (semantic β = 0.053, 95% CI [0.028, 0.078]; visual: β = 0.062, 95% CI [0.026, 0.098]; Fig. 4C). Taken together, the subicular signal exhibited the defining behaviour of a global comparator, in that it was greatest when memory afforded least match, diminished as accumulating experience rendered the present more predictable, and recovered as that experience receded in time.

## Discussion

Using ultra-high-field 7T fMRI during a continuous object recognition task, we tested whether the human hippocampus computes a non-associative, global, item-based mismatch signal, and if so in which subfield this computation is expressed in a behaviourally-grounded manner. Two dissociable signals predicted successful novelty detection: a univariate novelty response, largest in CA3 but most reliably predictive of detection success in CA1, and a multivariate global mismatch signal in the subiculum that predicted detection success independently of CA1. The subicular signal also propagated false alarms into later hits and scaled with the experienced similarity of past events. Together, these findings place CA1 and subicular comparator functions within a single computational account and broaden the hippocampal contribution to novelty detection beyond local, relational mismatch to include global, item-based matching. By delineating distinct hippocampal novelty signals that can bias decisions, this work provides a mechanistic scaffold for understanding, and ultimately targeting, aberrant novelty signalling in memory disorders.

First and foremost, our results extend the hippocampal comparator beyond its classical relational role. CA1 has long been identified as the site where expected and perceived information are compared, afforded by its direct entorhinal input connections alongside with indirect information fed through the DG and CA3 (*11*). Consistent with this anatomy, CA1 activity scales parametrically with relational mismatch (*22, 23, 28*), supporting accounts in which the hippocampus signals novelty through associative processing (*30, 31*). This relational emphasis reflects a broader view of the hippocampus as an associative system that binds items into contexts (*49–53*). However, a growing body of work also implicates the hippocampus in operations that do not require relational processing, including repetition suppression and old/new recognition effects (*15, 54*). The global mismatch signal we observed in this study falls on this non-associative, familiarity-like axis, classically attributed to the extra-hippocampal cortex (*55, 56*). In contrast, our results indicate that this signal is expressed in the subiculum, a core component of the hippocampal formation (*40*), which is consistent with intracranial evidence that the hippocampus supports both recollection and familiarity (*46*).

Building on this, here we propose that the subiculum implements a global-matching computation that complements the relational comparator in CA1. Global-matching models formalise recognition as the parallel comparison of a probe against all stored memory traces (*32, 33*). Such matching has traditionally been linked to familiarity and pattern-separated hippocampal representations have long been postulated to poorly suit this process (*35, 39*). This dependence on representational similarity, however, fits our observation that the subicular global mismatch signal scaled with the experienced similarity of prior events, in line with related work (*38*). Furthermore, the subiculum is anatomically well positioned for this global comparator role. As the principal output stage of the hippocampus, it pools convergent CA1 input (*41, 57*) and contains cells that respond selectively to novel objects (*58*). Framed within predictive coding, CA1 and the subiculum differ less in the type of computation than in its granularity (*41, 59*). While the CA1 novelty response might signal a local, relational prediction error, the subicular computation reflects a global, item-based prediction error that aggregates mismatch across accumulated experience. Furthermore, consistent with a role in the recognition decision itself, the behavioural influence of the subicular signal varied across individuals in a manner predictable from signal-detection measures. Participants with a more conservative criterion relied more strongly on the global mismatch signal when judging novelty, and those with higher sensitivity showed tighter coupling between this signal and subsequent retrieval success. Taken together, these results provide compelling evidence for the subicular contributions to a global mismatch signal for novelty detection.

Despite this evidence, several alternative interpretations warrant further consideration. First, novelty detection itself could differentiate subicular patterns, but the temporal resolution of fMRI cannot establish whether global mismatch is computed before or after the old/new decision. Nevertheless, it is important to note that this reverse-causality account predicts no relationship between first-encounter mismatch and future behaviour, whereas in our study a higher global-matching index both impaired novelty detection and increased the probability of successful retrieval on subsequent encounters. Second, univariate activation differences (i.e. novelty response) can reflect encoding differences, baseline shifts, or contextual prediction error (*15, 26, 60*), and disentangling pattern separation from univariate novelty signals remains challenging (*20, 21*). On the other hand, our multivariate measure is less vulnerable to these confounds, predicting novelty detection even after accounting for the univariate response. Third, stimulus attributes such as memorability, nameability, and recognisability predicted behaviour, indicating their overall contribution to the novelty detection process. Importantly, however, these stimulus factors did not correlate with the subicular global-matching index, indicating their minimal influence on the subicular computations. Together, these analyses support the interpretation that the subiculum implements computations consistent with the global matching process. Our conclusions are nonetheless constrained by stimulus scope, spatial resolution and task design. Since we only examined everyday objects embedded in naturalistic scenes, and because hippocampal engagement depends on goals, task demands and material (*22, 54, 61*), future work should further test other stimulus classes, including cut-out objects and verbal or abstract stimuli. Moreover, despite high-resolution 7T imaging and satisfactory Dice coefficients, we cannot fully exclude residual partial-volume blurring between CA1 and the adjacent subiculum and CA2. Therefore, methods such as sub-millimetre acquisitions could more precisely isolate each subfield’s contribution, although even under this limitation our findings would still implicate the hippocampus’ role in item-based novelty detection. Finally, whereas most memory studies use fewer than ~100 memoranda across one or two sessions with separated encoding and retrieval phases (*62*), our continuous design spanned five days and thousands of trials to approximate everyday experience (*63*), which may explain our ability to detect a global-matching signal in the subiculum. Nevertheless, future work using expanded time scales with larger sampling of the stimulus space will be required to deduce the definitive role that the human subiculum plays in novelty detection.

In conclusion, our findings raise several concrete questions: whether comparable global-matching signatures arise in the subiculum for other stimulus classes, how this signal is altered in ageing and patient populations (*7, 64*), and how such activity patterns can be integrated into formal global-matching models. More broadly, although hippocampal function has historically been framed in relational terms, our results suggest that this framework should be further expanded. Alongside the local, relational comparison attributed to CA1, the subiculum contributes a global, item-based mismatch signal that broadens the hippocampal role in novelty detection. Importantly, a clearer understanding of this subregional division of labour is likely to be consequential not only for basic accounts of hippocampal computation, but also for clinical contexts in which hippocampal circuitry is compromised (e.g., Alzheimer’s disease, temporal lobe epilepsy), where aberrant novelty signalling may contribute to cognitive symptoms and could offer more precise targets for intervention.

## Acknowledgements

We thank Peixin Yang and Yueting Su for their assistance in data acquisition. We also thank the staff at the Zhangjiang International Brain Imaging Centre (ZIC) for support during data collection, Ying-Hua Chu (MR Research Collaboration Team, Siemens Healthineers Ltd.) for assistance with protocol setup, and Matthew F. Glasser (Mallinckrodt Institute of Radiology, Washington University School of Medicine in St. Louis) and Essa Yacoub (Center for Magnetic Resonance Research, Department of Radiology, University of Minnesota) for their advice on optimising HCP-style data acquisition protocols.

D.V. discloses support for the research of this work from the Ministry of Science and Technology of China, STI2030 – Major Projects [grant number 2022ZD0207900]. J.A.Q received funding from the China Postdoctoral Science Foundation (2022M720818) to complete this work. All other authors declare no relevant funding. B.C.B. acknowledges support from the Canadian Institutes of Health Research, CIHR (FDN-154298, PJT-174995, PJT-206196, CIHR PJT-203761, CIHR PJT-191853), SickKids Foundation (NI17-039), Natural Sciences and Engineering Research Council (NSERC RGPIN-2025-05932), Azrieli Center for Autism Research of the Montreal Neurological Institute (ACAR), BrainCanada, FRQ-S, the Helmholtz International BigBrain Analytics and Learning Laboratory (Hiball), Healthy Brains and Healthy Lives (HBHL), Centre for Aging + Brain Health Innovation (CABHI), the Canada Research Chairs Program (CRC), and the Centre of Excellence in Epilepsy at the Neuro (CEEN).

## Author contributions

J.A.Q. and D.V. contributed to the conception and design of the work, while Y.W. and D.V. collected the data. The data were analysed by J.A.Q, K.Z., X.L., Y.W., J.F., J.D., B.C.B. and D.V. The first draft was written by J.A.Q. and D.V. All authors contributed to revision of the manuscript.

## Materials and Methods

### Participants

Twenty healthy young adult participants (14 females, 6 males; mean age = 24.56 years, SD = 2.42, range = 19–29) were recruited from the local university community according to predefined inclusion criteria. All participants were right-handed native Mandarin speakers with normal or corrected-to-normal vision and reported no known history of neurological or psychiatric conditions and no contraindications to MRI. The sample size was chosen in line with recent fMRI experiments employing dense sampling procedures, which prioritise per-participant data quantity over cohort size to support robust individual-level inference (*65, 66*). After a detailed explanation of the study aims and the multi-session protocol, participants provided written informed consent and were compensated at 100 RMB per hour. All procedures adhered to the Declaration of Helsinki – Ethical Principles for Medical Research Involving Human Participants and were approved by the local institutional review board (protocol AF/SC18/20210604).

### Methods details

#### Experimental procedures

Each participant completed six visits at the Zhangjiang International Brain Imaging Centre (ZIC), Fudan University. The first visit was dedicated to 3T anatomical imaging (T1w, T2w) and resting-state fMRI. The remaining five visits, typically scheduled on consecutive weekdays (Monday–Friday) at approximately the same time of the day, were dedicated to 7T task fMRI (median interval between successive 7T sessions = 23.8 h). At every 7T visit, participants performed a continuous object recognition task and a functional localiser (the latter not analysed here).

### Stimuli

Stimuli for the continuous object recognition task (*65*) were drawn from the THINGS database (*43*), which comprises 22,256 high-quality naturalistic object images spanning 1,854 diverse object concepts (e.g., aardvark, airplane, shoes). Each image depicts a single object embedded in a natural scene context (e.g., a bicycle leaning against a wall; Fig. 1A). The image set is accompanied by extensive annotations including dimensions that explain a substantial fraction of variance in human similarity judgements (*47, 67*), making it well suited to studying novelty and memory within a continuous recognition design.

### Continuous object recognition task

On each trial, a single object image was presented on a uniform grey background while participants maintained central fixation on a small transparent red dot. Participants indicated by button press whether the image was *new* (never previously seen across any session) or *old* (presented during the current or any earlier session). Each trial lasted 4 s, comprising 3 s of image presentation followed by a 1 s inter-trial blank. Each run included 75 trials in total: 64 image trials and 11 fixation-only blank trials, with three blanks placed at the start, four at the end and four pseudo-randomly between image trials, such that consecutive image trials appeared in stretches of 9-14. At the end of each run, participants received feedback indicating the number of responses recorded.

Across the five 7T sessions, each participant completed 60 task runs in total (12 runs per session). With the aim of maximising wide coverage of the THINGS taxonomy, we selected 22,256 images spanning all 1,854 concepts (13 images for 8 concepts and 12 images for all other), drawn as the first 12 (or 13) entries in each concept’s image list. The only a priori, content-based exclusion was the removal of images depicting threat objects (e.g., firearms pointed directly at the viewer); no further concept- or image-level exclusions were applied. The THINGS image pool was partitioned into 176 shared images (presented to every participant) and 22,080 unique images distributed evenly across participants (1,104 unique images per participant). Each participant therefore viewed 1,280 distinct images (176 shared + 1,104 unique), each presented three times, resulting in 3,840 image trials per participant. On average, each concept was sampled by 1.35 unique exemplars per participant (SD = 0.02).

Following prior work (*65*), the sequence of 3,840 trial slots, including the temporal lag structure between successive presentations of the same image, was generated from a mixed distribution (60% von Mises, κ = 10; 40% uniform) and applied identically across participants. Within this fixed template, the 176 shared-image slots were occupied by the same images for every participant, whereas the remaining slots were filled with each participant’s unique image set. As a result, the temporal structure of repetitions (the lag distribution between the first, second and third presentations of each image) was matched across participants while the underlying images differed. This procedure also ensured that participants continued to encounter novel images including in the final session.

Due to technical issues encountered during data acquisition, two runs had to be repeated, affecting two participants. Consequently, a small subset of images was presented more than three times to these individuals. To preserve the intended novelty structure of the task, images encountered in repeated runs, together with their subsequent presentations, were excluded from all analyses.

### MRI data acquisition

Neuroimaging data acquisition followed Human Connectome Project (HCP) guidelines (*68*). Anatomical and resting-state fMRI scans were acquired on a 3T Siemens Magnetom Prisma (Siemens, Erlangen, Germany) with a 32-channel head coil, using sequences adapted from HCP-Lifespan (*69*). Task fMRI was acquired on a 7T Siemens Magnetom Terra fitted with a 1-channel transmit / 32-channel receive (1Tx/32Rx) head coil (Nova Medical, Wilmington, MA, USA), using harmonised sequences from the UK-7T network (*70*). Fully compatible with the HCP preprocessing pipelines (*71*), this dual-scanner design combined the wide field-of-view and low geometric distortion of 3T anatomical imaging with the high spatial resolution and signal-to-noise ratio afforded by 7T for task-evoked BOLD activity.

At 3T, whole-brain anatomical imaging comprised a T1-weighted MPRAGE sequence (0.8 mm isotropic; TR = 2,500 ms; TE = 2.22 ms; TI = 1,000 ms; flip angle = 8°; bandwidth = 220 Hz/pixel; in-plane acceleration [iPAT] = 2; acquisition time [TA] = 6 min 40 s) and a T2-weighted SPACE sequence (0.8 mm isotropic; TR = 3,200 ms; TE = 563 ms; bandwidth = 745 Hz/pixel; iPAT = 2; TA = 5 min 57 s). Resting-state fMRI (eyes open, fixating a central crosshair) used a multiband gradient-recalled echo-planar imaging (MB-GRE-EPI) sequence (2.0 mm isotropic; TR = 800 ms; TE = 37 ms; flip angle = 52°; bandwidth = 2,290 Hz/pixel; multiband factor = 8; 488 volumes; TA = 6 min 40 s), acquired with both anterior-posterior (AP) and posterior-anterior (PA) phase-encoding runs to enable post hoc distortion correction. Additional spin-echo and dual-echo gradient field maps were collected for geometric distortion correction.

Task fMRI at 7T used a comparable MB-GRE-EPI sequence optimised for spatial resolution (1.5 mm isotropic; TR = 1,500 ms; TE = 25 ms; flip angle = 65°; multiband factor = 4; phase-encoding direction = AP; TA = 8 min 40 s per run). A matched pair of dual-echo gradient field maps was acquired before every four task runs to correct susceptibility-induced distortions. Visual stimuli were back-projected onto a screen at the rear end of the bore using an MRI-compatible PROPixx projector (VPixx Technologies, Canada) and viewed through a mirror mounted on the head coil (viewing distance = 202.5 cm; display resolution = 1920 × 1080; refresh rate = 120 Hz; image resolution = 800 × 800 pixels; visual angle ≈ 6°). Behavioural responses were collected with an MRI-compatible four-button response box.

### MRI data preprocessing

All neuroimaging data were processed with the HCP minimal preprocessing pipelines (*71*), as implemented in the Quantitative Neuroimaging Environment & Toolbox (QuNex, version 0.99.1) (*72*). Structural images were corrected for gradient non-linearity and aligned to MNI152 space; tissue segmentation and cortical surface reconstruction were performed in FreeSurfer (Version 6.0.7.1) (*73*). Functional data were corrected for gradient and EPI susceptibility distortions and for head motion, and registered to the MNI152 space. Except for the removal of spatially specific temporal artefacts using the ICA+FIX approach (*74*), no additional temporal filtering, detrending or spatial smoothing was applied, beyond the native 2 mm FWHM. In order to align the volumetric BOLD signal timeseries with the timing of trials in the task design, preprocessed timeseries were up-sampled to 1 s intervals using the *tseriesinterp* function from the tool collection *knkutils*.

### Trial-wise neural response estimation

Trial-wise BOLD response amplitudes for the task were estimated with GLMsingle (*75*), which augments a standard GLM analysis with important optimisation steps. Since our analyses focused on changes in response magnitude across repeated presentations of the same image, we used the TYPEB beta estimates (b1), which incorporate per-voxel selection of an optimal hemodynamic response function (HRF) from a library of 20 candidates but omit the GLMdenoise and ridge-regression steps, in line with recent applications of GLMsingle (*76*). Trial onsets were rounded to the nearest upsampled volume. Amplitudes were converted to percent BOLD change but were otherwise left unscaled in order to preserve unbiased estimation of repetition- and novelty-related responses.

### Hippocampal segmentation and surface mapping

Hippocampal subfields were segmented and participant-specific surfaces were generated using HippUnfold (version 1.5.2) (*44*), run as a Singularity container. We segmented the AC-PC-aligned, distortion-corrected T1w images from the structural HCP preprocessing pipeline, with the output surface density set to 0.5 mm (Dice coefficients all > 0.7, minimum 0.72; cross-hemisphere mean 0.82, SD = 0.012). The resulting surfaces were warped to MNI152 space using each participant’s forward and inverse transforms. We then sampled single-trial beta estimates onto the MNI152-aligned surfaces with ribbon-constrained averaging across the inner, midthickness and outer cortical depths. The resulting per-vertex data were downsampled to 2 mm resolution and stored as HDF5 files to streamline downstream handling. Regions of interest (ROIs) were defined on the multihist7 subfield atlas (*77*) distributed with HippUnfold. Owing to the limited resolution of our acquisition, results for CA4 are omitted, and the final set of hippocampal subfields included in this study covered CA1-CA3, subiculum and DG.

### Quantification and statistical analysis

#### Signal detection measures

In order to characterise individual differences in memory performance, we computed standard signal detection measures. Sensitivity was quantified as d′:

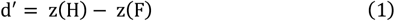

and the decision criterion (response bias) as c:

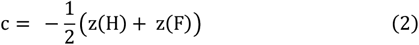

where z(p), p ∈ [0,1] is the inverse of the standard normal (Gaussian) cumulative distribution function, and H and F denote the hit and false-alarm rates, respectively, which were calculated with log-linear transformations. Negative values of c indicate a liberal response bias (a greater tendency to endorse an image as old), positive values a conservative bias, with a neutral point of zero.

### Behavioural and neural data quality analyses

We assessed the behavioural data for signs of fatigue or waning motivation, which might manifest as declining response rates over the course of the experiment. For this, we modelled the probability of responding on a given trial as a function of running trial number in a logistic regression model, and tested whether average run-wise reaction time changed systematically across the experiment in a Gaussian regression model.

Since the number of items to be remembered greatly exceeded that of conventional recognition experiments, we further examined whether performance changed over time. Making full use of the dataset, including the final runs, requires that participants could still discriminate old from new images by the end of the experiment. We therefore submitted session-wise response bias and d′ to Gaussian regression models to characterise how each measure evolved across sessions. Critically, d′ remained above chance in every session, including the last.

Neural data quality was quantified in two ways. First, we computed the TYPEA beta estimate (b0) per participant and ROI, which captures the explained variance (R2) of the ON–OFF model. Second, we computed temporal signal-to-noise ratio (tSNR; the mean divided by the standard deviation of the timeseries) averaged across all runs. Each measure was analysed with a Bayesian linear model, in which we extracted marginal means for each subfield and computed contrasts comparing each subfield against the unweighted mean of the others, resulting in a posterior median and 95% credible interval for each comparison. All comparisons are provided in the Supplementary Information (Table S2 and S3). We then calculated the Bayesian Pearson correlation between these neural data quality metrics and the participant-level slopes in the main hierarchical regression models (see below) to ensure that neural data quality did not affect our inference with regard to predicting behaviour based on the magnitude of the novelty response and global mismatch. None of the correlations showed robust associations.

### Univariate novelty response analysis

Our first main analysis characterised the classical novelty response across hippocampal subfields, modelling the difference in BOLD signal between the first and second encounter of an image (1^st^ > 2^nd^) in Gaussian regression models. In order to avoid multicollinearity, each subfield was modelled separately throughout the remaining analyses unless explicitly stated otherwise. We then tested whether the magnitude of the novelty response was diagnostic of trial-wise novelty detection success (i.e. correctly judging a new image as new) in a series of logistic regression models.

### Multivariate global mismatch analysis

For each ROI we extracted trial-wise neural activation vectors and computed a trial-by-trial neural similarity matrix as the Fisher-z-transformed Pearson correlation between activation patterns. For every trial, we defined the neural global matching index as the maximum correlation between the current trial and all previously presented trials, following classical global matching formulations (*32, 33*). On this logic, low values of the index constitute a global mismatch, which we hypothesised would contribute both to successful novelty detection and to the propagation of false alarms. Trials from the same run as the target were excluded to guard against temporal autocorrelation in the BOLD signal (*76*). The index was tested as a predictor of accuracy on first-encounter trials (novelty detection) separately for every subfield and without privileging any sub-field a priori. Additionally, we used this index to model second and third encounters (retrieval).

In a control analysis we tested whether image-level memorability, nameability and recognisability, rich annotations available for the THINGS stimulus set (48), predicted the global matching index in Gaussian regression models, addressing whether more memorable or nameable images elicit systematically higher indices (a potential confound for our primary analysis).

### Semantic and visual similarity analysis

Beyond image-level features, THINGS annotations include 66 dimensions per image that capture the majority of variance in human similarity judgements across concepts and images (*47*). Following an established protocol (*48*), we classified each dimension as reflecting semantic information (categorical information without reference to visual properties; e.g. electronics/technology), visual information (shape or colour; e.g. black) or a mixture of the two (e.g. paper-related/flat). Treating the semantic and visual dimensions as separate embedding vectors, we computed image-to-image correlation matrices using the same logic as for the neural global matching index, defining semantic and visual similarity for each trial as the maximum correlation with all images presented up to that point. The semantic and visual representational structures were visualised with 2D t-SNE maps coloured by their highest-level category.

We then used these similarity metrics to predict both novelty detection success and the neural global matching index, testing the prediction that recently encountered similar images raise global matching (reduce mismatch) and thereby impair detection. As a further control analysis, we extracted embeddings for each image from a pretrained CLIP model (*78*) and repeated these analyses using CLIP-based similarity in place of the similarity derived from the 66 dimensions.

### Bayesian statistical modelling

Unless stated otherwise, all main analyses were conducted at the trial/image level, allowing us to model trial-unique representations and responses, to resolve the effects of temporal lag at high resolution, and to contrast competing hypotheses directly through control analyses. Running trial number was included as a covariate to account for the changing old/new ratio over the course of the experiment and for the fact that maximum correlation coefficients increase mechanically as the pool of previously seen images grows.

Our main hypotheses were tested with Bayesian hierarchical regression models (*79*). All models included full participant-level random effects (random intercepts and slopes) and were estimated with five Markov chains of 6,000 iterations each (3,000 warm-up, 3,000 sampling). Convergence was confirmed by 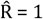 for all parameters, and the adequacy of each response distribution was evaluated in posterior predictive checks. Continuous predictors were standardised to mean = 0 and SD = 0.5; for Gaussian models the dependent variable was additionally scaled to mean = 0 and SD = 1. Following previous work (*76*), temporal lags (in minutes) were logtransformed prior to scaling.

For logistic (Bernoulli), Gaussian and lognormal reaction-time models alike, weakly informative priors were used throughout, following standard recommendations for hierarchical GLMs. For Gaussian models, predictors were mean-centred so the intercept was fixed at zero. The complete prior specifications for every model are tabulated in the accompanying repository. For every model we report posteriors means of the regression coefficients with 95% credible intervals (CIs), treating an effect as robust only when its 95% CI excluded zero. Bayesian versions of *t*-tests and bivariate Pearson correlations were computed with the R package *BayesFactor* using its default prior.

## Supplementary Materials

**Fig. S1.**
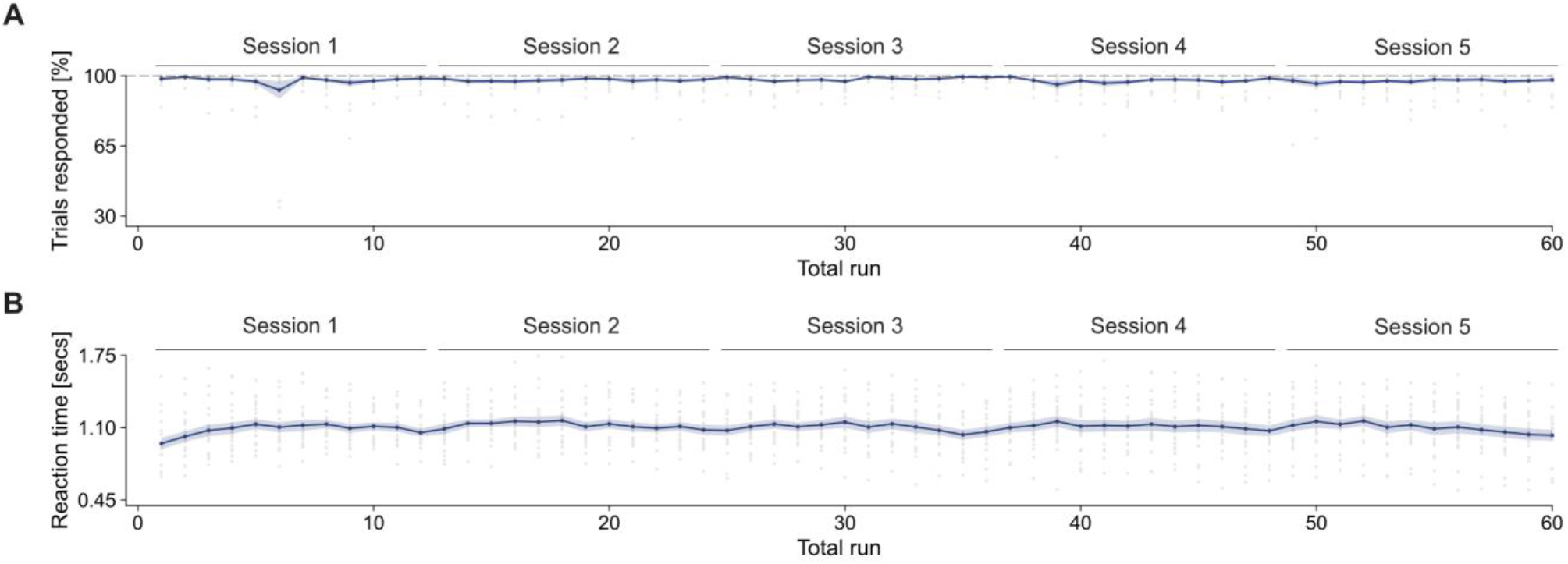
Response rates and reaction times over the course of the experiment. **(A)** Run-level response rates, which remained high (82–100%) with no decline across the experiment. **(B)** Similarly, run-averaged reaction times were stable at ~1.1 s. Both metrics highlight consistent task engagement throughout the experiment. Grey dots denote participant-level data, while the coloured lines and ribbons show group-level mean ± standard error of the mean (SEM).

**Fig. S2.**
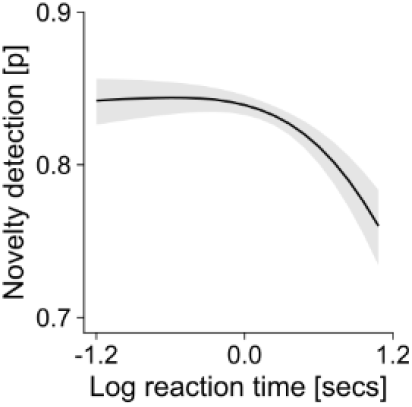
Reaction time predicts novelty detection. Shorter (log) reaction times predicted more accurate novelty detection (β = −0.55, 95 % CI [-0.96, −0.16]), supporting the use of decision speed as a proxy for decision confidence (see main-text Figure 3B for the complementary link between the subicular matching index and response speed). Marginal averaged predictions are shown with 95% CI.

**Fig. S3.**
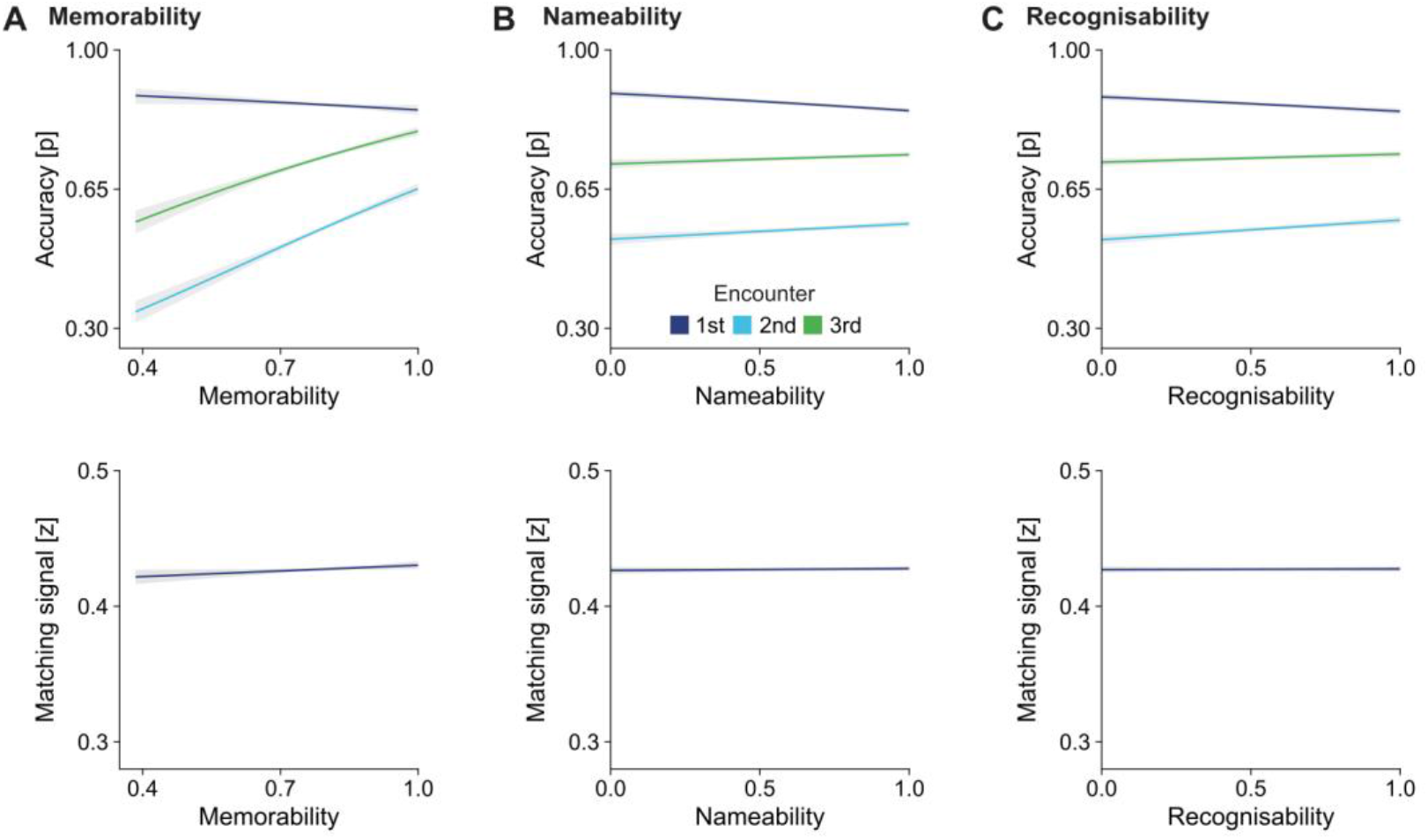
Effects of memorability, nameability and recognisability on trial accuracy and global mismatch. **(A)** Higher memorability decreased accuracy for the 1st encounter (−0.0091, 95% CI [−0.017, −0.0013]), but increased for the 2nd encounter (0.078, 95% CI [0.068, 0.089]) and also increased for the 3rd encounter (0.054, 95 % CI [0.046, 0.062]). There was no relationship with global matching in the subiculum (β = 0.02, 95 % CI [−0.00077, 0.041]). **(B)** Higher nameability decreased accuracy for the 1st encounter (−0.026, 95 % CI [−0.035, −0.017]), increased for the 2nd encounter (0.022, 95 % CI [0.011, 0.032]) and also increased for the 3rd encounter (0.013, 95 % CI [0.0047, 0.022]). There was no relationship with global matching in the subiculum (β = 0.0076, 95 % CI [−0.013, 0.029]). **(C)** Higher recognisability decreased accuracy for the 1st encounter (−0.02, 95 % CI [−0.029, −0.012]), increased for 2nd encounter (0.028, 95 % CI [0.017, 0.038]) and also increased for the 3rd encounter (0.011, 95 % CI [0.0026, 0.02]). There was no relationship with global matching in the subiculum (β = −0.0036, 95 % CI [−0.031, 0.02]). Marginal averaged predictions as a function of an image’s feature; shaded areas are 95% CI, with separate slopes for the first (orange), second (green) and third (blue) encounters to visualise the interaction, when predicting accuracy.

**Fig. S4.**
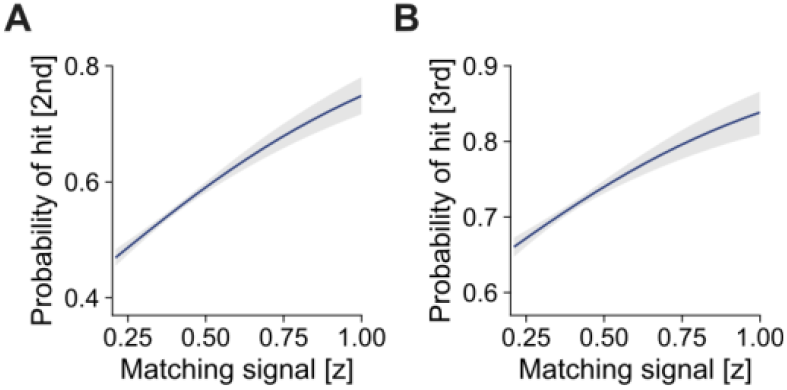
Propagation of the first-encounter subicular matching signal into later behaviour. A high global match on the first encounter predicts subsequent hits on the 2^nd^ **(A)** and 3^rd^ **(B)** encounters, the diagnostic signature of a global-matching comparator. Marginal averaged predictions are shown with 95% CI

**Fig. S5.**
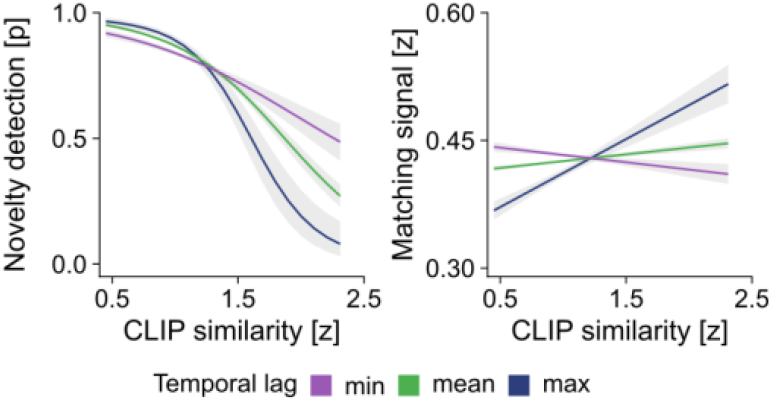
Replication with CLIP-based similarity. The semantic and visual similarity analyses of Figure 4 was repeated using image embeddings from a pretrained CLIP model (ViT-L/14@336px; Radford et al., 2021) in place of the 66 THINGS similarity dimensions. On the behavioural level, this analysis revealed that when CLIP similarity was high, the probability of novelty detection success was low (β = −1.2, 95 % CI [−1.4, −1.1]). This again was modulated by the log-transformed lag in minutes between the current trial and the encounter of the most similar image (β = 0.53 (95 % CI [0.28, 0.79]). In line with the general results, CLIP similarity was positively correlated with the neural matching signal (β = 0.059, 95 % CI [0.032, 0.083]) and the influence of similarity was again reduced at longer lags (β = −0.15, 95 % CI [−0.19, −0.11]). The interaction is visualised with minimum lag (blue), average lag (green) and maximum lag (purple). Marginal averaged predictions are shown with 95% CI.

**Fig. S6.**
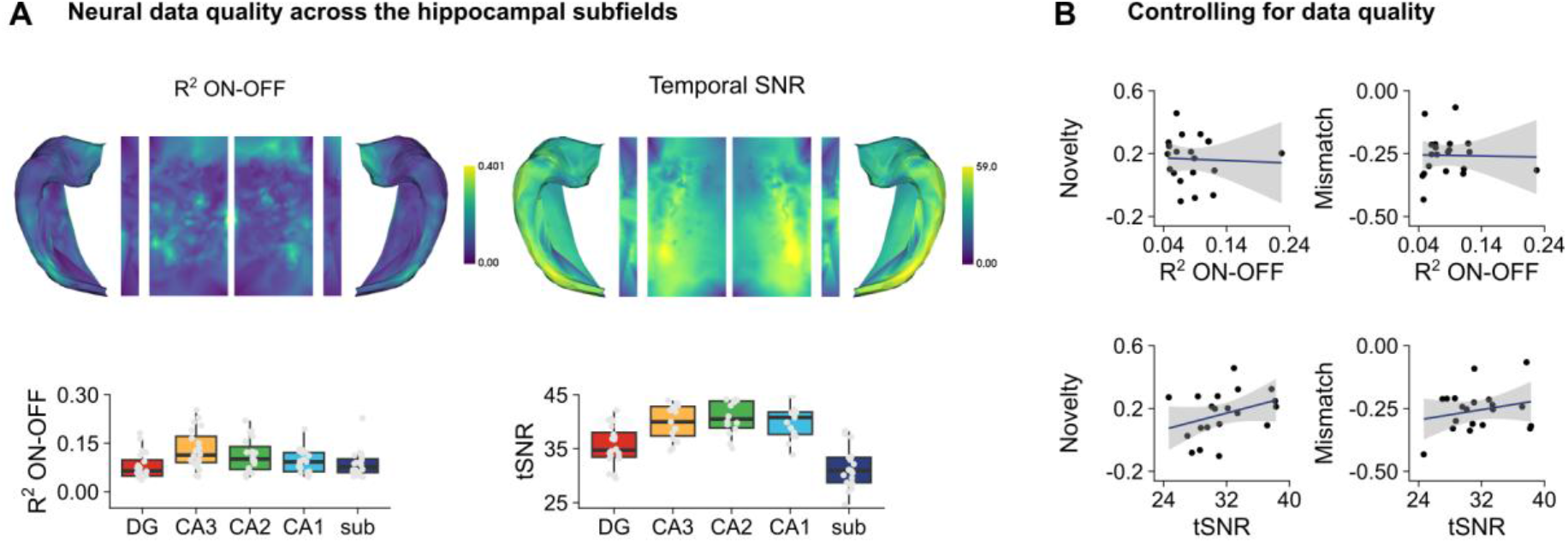
Neural data quality across hippocampal subfields. **(A)** Per-participant neural data quality for each subfield: explained variance (R^2^) of the ON–OFF model (TYPEA b0 estimate) and temporal signal-to-noise ratio (tSNR), averaged across runs. Average metrics per subfield per person are shown as boxplots (for comparisons see Table S2 & S3). **(B)** In order to investigate whether individual differences in neural data quality in the subiculum affected our results, we correlated each metric (R^2^ and tSNR) with the participant slopes derived from the model of the novelty response (left column) and global mismatch (right column) predicting novelty detection, None of the correlations showed any evidence for an association: novelty & R^2^, r(18) = −.045, BF_10_ = 0.48, novelty & tSNR, r(18) = .345, BF_10_ = 1.14, mismatch & R^2^, r(18) = −.023, BF_10_ = 0.48, and finally mismatch & tSNR, r(18) = .235, BF_10_ = 0.7.

**Table S1.** Classification of the 66 dimensions.

| **Semantic** | **Visual** | **Mixed** |
| --- | --- | --- |
| metallic/artificial | circular / round | wood-related/brown |
| food-related | white | colorful / playful |
| animal-related | red | paper-related / flat |
| textile | long / thin | tools-related / handheld / elongated |
| plant-related | black | coarse-scale pattern / many things |
| house-related/furnishing-related | tubular | stick-shaped / container |
| valuable/precious | grid-related / grating-related | thin / flat / wrapping |
| transportation-/movement-related | repetitive / spiky | cylindrical / conical / cushioning |
| body-/people-related | spherical / voluminous |  |
| electronics / technology | string-related / stringy / curved |  |
| outdoors | transparent / shiny / crystalline |  |
| hobby-related / game-related / playing-related | sand colored |  |
| fluid-related / drink-related | green |  |
| water-related | yellow |  |
| oriented / many things | upright / elongated and volumous |  |
| powdery / earth-related / waste-related | pointed / spiky |  |
| weapon-related / war-related / dangerous | orange |  |
| household-related | fine-grained pattern |  |
| feminine (stereotypical) |  |  |
| body part-related |  |  |
| music-related / hearing-related / hobby-related / loud |  |  |
| construction-related / craftsmanship-related / housework-related |  |  |
| seating / standing / lying-related |  |  |
| flying-related / sky-related |  |  |
| bug-related / non-mammalian / disgusting |  |  |
| bathroom-related / wetness-related |  |  |
| heat-related / fire-related / light-related |  |  |
| beams-related / mesh-related |  |  |
| foot-related / walking-related |  |  |
| box-related / container |  |  |
| head-related |  |  |
| child-related / toy-related / cute |  |  |
| farm-related / historical |  |  |
| seeing-related |  |  |
| medicine-related / health-related |  |  |
| sweet / dessert-related |  |  |
| coldness-related / winter-related |  |  |
| measurement-related / numbers-related |  |  |
| fluffy / soft |  |  |
| masculine (stereotypical) |  |  |

**Table S2.** Comparing each subfield’s R2 with the average of all other via marginal means.

| Subfield | Median | 2.5 % | 97.5% |
| --- | --- | --- | --- |
| CA1 | -0.010 | -0.025 | 0.0037 |
| CA2 | 0.025 | 0.011 | 0.0390 |
| CA3 | 0.035 | 0.021 | 0.0490 |
| DG | -0.028 | -0.042 | -0.0140 |
| sub | -0.021 | -0.036 | -0.0075 |

**Table S3.** Comparing each subfield’s tSNR with the average of all other via marginal means.

| Subfield | Median | 2.5% | 97.5% |
| --- | --- | --- | --- |
| CA1 | 3.9 | 3.1 | 4.6 |
| CA2 | 5.1 | 4.3 | 5.9 |
| CA3 | 3.9 | 3.1 | 4.7 |
| DG | -3.9 | -4.7 | -3.2 |
| sub | -8.9 | -9.7 | -8.2 |

